# Comparative lipidomic profiling of the livestock pathogens *Trypanosoma congolense* and *Trypanosoma brucei*

**DOI:** 10.64898/2026.04.21.719795

**Authors:** Emily A. Dickie, Stefan K. Weidt, Jennifer Haggarty, Gavin Blackburn, Mary K. Doherty, Ryan Ritchie, Edith Paxton, Simon Young, Phillip D. Whitfield, Terry K. Smith, Liam J. Morrison, Michael P. Barrett, Pieter C. Steketee

**Affiliations:** School of Infection and Immunity, University of Glasgow, Glasgow, United Kingdom; Shared Research Facilities (SRF), MVLS, University of Glasgow, Glasgow, United Kingdom; Division of Biomedical Sciences, University of the Highlands and Islands, Centre for Health Sciences, Inverness, United Kingdom; The Roslin Institute, Royal (Dick) School of Veterinary Studies, University of Edinburgh, Midlothian, United Kingdom; Schools of Chemistry and Biology, Biomedical Sciences Research Complex, North Haugh, University of St Andrews, St Andrews, United Kingdom

## Abstract

African Animal Trypanosomosis (AAT) is a disease affecting domestic animals, in particular cattle, in sub-Saharan Africa, resulting in billion-dollar losses annually. New drugs to combat and control AAT are urgently required, yet few treatment candidates are currently on the horizon. This can be attributed, in part, to the relative challenges associated with culturing the clinically relevant parasite species in a laboratory environment. Particularly, effective culture of bloodstream form *Trypanosoma congolense*, the trypanosome species responsible for a large proportion of AAT disease in cattle, requires the use of goat serum, whilst *T. brucei* is typically cultured in FBS-supplemented culture. This constrains *in vitro* studies on biology, especially comparative analyses between AAT-causing species. The differing serum supplementation requirements of these two trypanosome species point to metabolic distinctions, which may be important considerations in developing experimental systems to enable the identification and design of novel, pan-species therapies. In this study, untargeted LC-MS lipidomics analyses were conducted to determine the relative lipidomic profiles of *T. congolense* and *T. brucei* bloodstream form parasites. Employing a new media formulation that permits effective *in vitro* culture of both species, it was possible to establish that their global lipidomic profiles are distinct. Notably, *T. congolense* exhibits a relatively low abundance of ether phospholipids compared to *T. brucei*, whilst also possessing an enrichment of long-chain polyunsaturated fatty acids (PUFAs). These observations indicate that there are significant differences in the ways these parasites synthesise and remodel their lipid complement, highlighting an evolutionary divergence between the species that likely carries implications for host-pathogen interactions as well as trypanosome membrane biology. Furthermore, this study demonstrates that fine-tuning fatty acid supplementation may aid in optimising a universal medium suited for multiple species of AAT parasites.

**Summary:** Multiple species of protozoan parasites can cause African Animal Trypanosomosis (AAT) in livestock and other animals. However, AAT research has largely centred on a single species, *Trypanosoma brucei*, partially due to the comparative difficulties in sustaining the other economically important parasite species - *Trypanosoma congolense* and *Trypanosoma vivax* - in laboratory culture. In this work, we aimed to determine whether distinctions in use of lipids between *T. brucei* and *T. congolense* explains their differing *in vitro* culture requirements. Using a newly designed media formulation, it was possible to culture mammalian-infective forms of both parasite species under identical conditions, enabling direct comparison of their lipidome - a complete inventory of the different fats and lipids the cells contain. We demonstrate that the *T. congolense* lipidome significantly differs from that of *T. brucei*, and that *T. congolense* shows a preference for longer, more unsaturated lipids. These differences are likely to underlie species-specific differences observed during host infections. Furthermore, our work demonstrates that understanding the lipid biology of protozoan parasites aids in optimisation of laboratory culturing conditions, thereby facilitating further research into these understudied pathogens, including the development of new therapies.

## Introduction

African Animal Trypanosomosis (AAT) is a parasitic disease that poses a significant risk to the health of livestock and other animals in sub-Saharan Africa. The disease is primarily caused by three species of African trypanosome; *Trypanosoma congolense*, *T. vivax* and *T. brucei* [1]. It is estimated AAT causes in excess of three million cattle deaths per year, with up to 90 million at risk of disease [2]. New treatment options to treat the disease are sorely needed. Historically, AAT has been largely neglected by drug development endeavours, although this has changed recently with the ongoing development of benzoxaborole compounds as potential novel trypanocides [3,4]. Consequently, no new drugs for AAT have been introduced in over 50 years, leading to increasing reports of parasite drug resistance to the two main chemotherapeutics (isometamidium chloride and diminazene aceturate) [2]. Establishing control of AAT infection is critical to protect economic interests (estimated economic losses to AAT amount to US$ 4.5 billion [5]) and to remove a significant barrier to the development of sustainable agriculture in the affected areas.

An impediment to developing drug treatments for AAT has been the relative difficulty of studying the different causative parasite species in a laboratory setting. Although *T. brucei* has been widely studied, in part due to sub-species of *T. brucei* (*T. brucei gambiense* and *T. brucei rhodesiense*) being responsible for HAT, it was only recently that mammalian-infective bloodstream form (BSF) *T. congolense* became more widely studied under *in vitro* conditions [6–8], and there is still no established method for the *in vitro* culture of BSF *T. vivax*. Furthermore, tools for the genetic manipulation of *T. congolense* have only recently become available, enabling the functional study of this species on a molecular level [6].

Whilst *in vitro* cultivation of *T. brucei* has been a laboratory mainstay for many decades [9], culture of *T. congolense* has proved far more challenging, with a key difference being a requirement for fresh goat serum (GS) supplementation of culture medium, in contrast to the more commonly used and inexpensive foetal bovine serum (FBS) for *T. brucei*, the latter in common with most other eukaryotic *in vitro* culture models [7,10]. The use of GS as serum supplement significantly hinders the ability to carry out direct comparative analyses of *T. congolense* and *T. brucei*, including their co-culture. It also raises a key question - how do these sera differ biochemically, and what are the key components *T. congolense* requires from GS that FBS cannot provide?

In previous work, we generated a comprehensive overview of the *T. congolense* metabolome [8], including a comparative analysis with *T. brucei*. Of particular note were differential responses to inhibition of fatty acid (FA) synthesis, with *T. congolense* exhibiting significantly reduced sensitivity to an inhibitor of acetyl-CoA synthetase and Orlistat, a known inhibitor of, amongst other targets, FA synthase [8,11]. These data suggest profound differences in FA biosynthesis, and likely, composition, between *T. brucei* and *T. congolense*. However, these studies were still carried out with the caveat that both species were cultured in different sera, the sole source of lipids under *in vitro* conditions.

In this study, we sought to analyse the metabolic signatures of the sera used to culture trypanosomes, and the BSF-stage parasites themselves, in order to generate a comparative analysis of lipid metabolism in *T. congolense* and *T. brucei* BSFs. Employing a new media formulation that supports the growth of both *T. brucei* and *T. congolense* BSFs, it was possible to establish that the lipidome of *T. congolense* is distinct from that of *T. brucei*, even when both species are subjected to the same *in vitro* media conditions. This points to a fundamental divergence in their lipid metabolism, which could have key implications for drug development and host-pathogen interactions. Targeted GC-MS analysis indicates that this divergence extends to the fatty acid content. Overall, results indicate it may be possible to fine-tune fatty acid provision to permit further culturing and/or co-culturing options for these trypanosome species, giving us greater insight into evolutionary divergences in lipid metabolism in these closely related pathogen species.

## Results

### Trypanosome growth *in vitro* is heavily influenced by serum source

Previous studies have emphasised serum source as an important factor for the successful cultivation of animal trypanosomes *in vitro* [7,9]. Notably, *in vitro* medium formulations described for *T. congolense* contain a goat serum supplement (20%), in contrast to *T. brucei,* which is typically supplemented with 10% FBS [9]. To date, no biological reason for this discrepancy has been validated. To characterise the viability of *T. congolense* and *T. brucei* in differing serum supplements, growth curves were carried out in HMI-11 (*T. brucei*) or HMI-93 (*T. congolense*) supplemented with 10% and 20% v/v of different animal sera (Fig 1).

**Figure 1:**
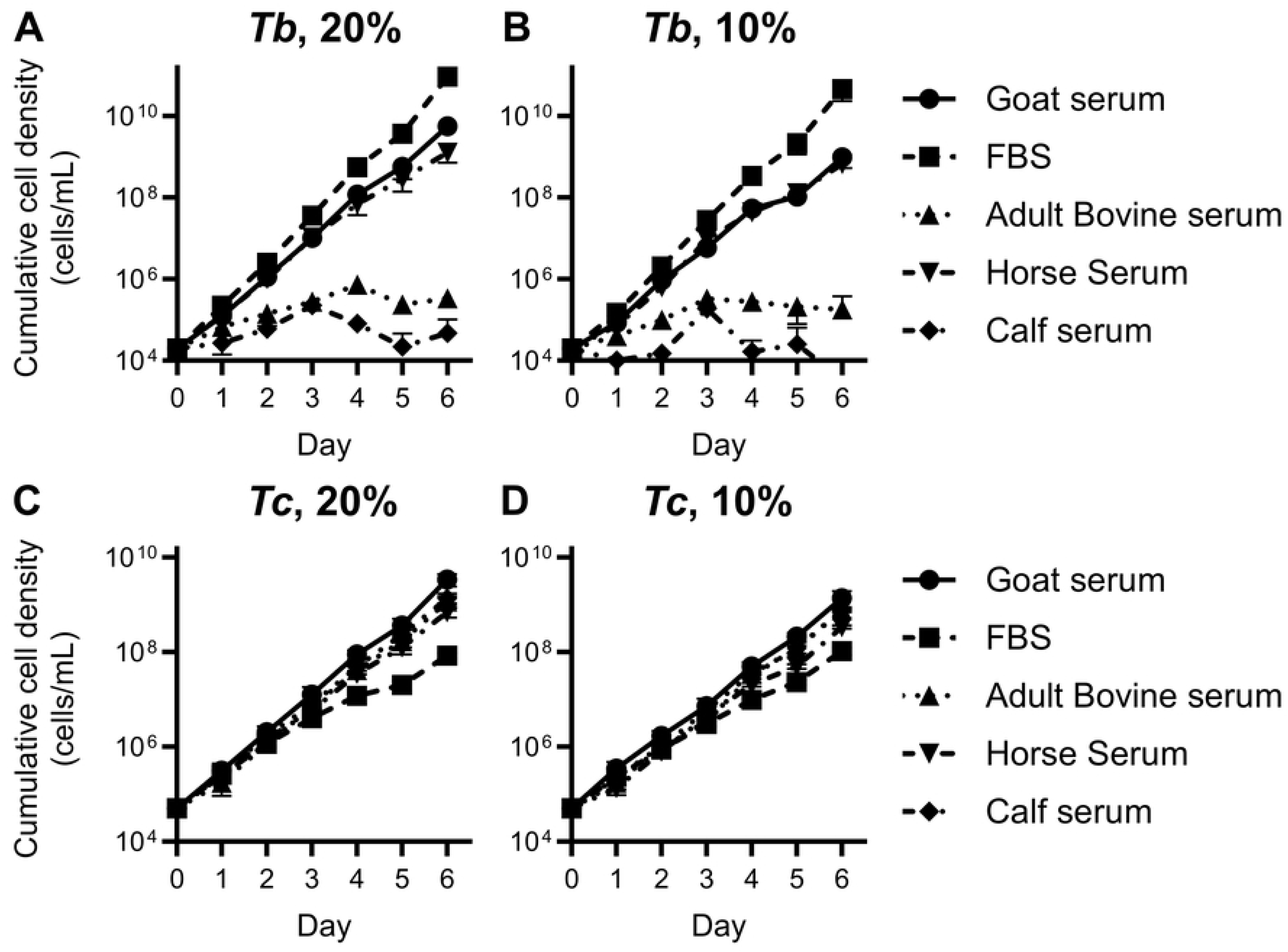
Effect of serum supplement on trypanosome *in vitro* viability. Cultures of *T. brucei* (A & B) and *T. congolense* (C & D) were seeded and cell density counted by haemocytometer every 24 hr in the presence of 20% (A & C) or 10% (B & D) serum. A total of five sera were tested: goat serum (GS), foetal bovine serum (FBS), adult bovine serum (ABS), horse serum (HS) and calf serum (CS). Cells were passaged every 48 hr, and cumulative cell density over a period of 6 days is shown. Doubling times were calculated and statistically compared (BH-corrected *t*-test) to the relevant control (10% FBS and 20% GS for *T. brucei* and *T. congolense*, respectively), and are presented in Fig S1.

As expected, *T. brucei* grows best in FBS, supplemented at either 20% or 10% (Fig 1A & 1B; doubling times shown in Fig S1). Cells remained viable in goat serum and horse serum, albeit with a reduced doubling time (Fig S1). Adult bovine serum and calf serum did not support *T. brucei* culture, with no cell division observed after 2-3 days of culture (doubling times could therefore not be calculated for these samples).

In contrast, *T. congolense* was, in general, viable in all sera tested (Fig 1C & 1D; doubling times shown in Fig S1). However, long-term culture in FBS was not successful. It was also determined that reduction of any serum to 10% (instead of 20%) led to a reduction in *T. congolense* doubling time (Fig S1).

The function of cell culture serum is to provide additional (undefined) nutrients vital to supporting the growth of cells *in vitro*. Serum contains sugars, such as glucose, amino acids and micronutrients such as inorganic ions and vitamins, which are supplied to considerable excess in basal media formulations, including IMDM, which is a main component of HMI-11 and HMI-93 [7,9]. Metabolomics analysis to assess potential differences in the small, polar molecule complement of FBS and GS revealed many differences in polar metabolites (Fig 2A). Of the 482 metabolites detected, 285 were significantly different (BH-corrected *t*-test; S1 Table), of which the top 50 are shown in Fig 2A, and biologically relevant metabolites shown in Table 1. These metabolites were mainly annotated as amino acids, such as L-aspartate (Log_2_FC, GS/FBS: -2.81), L-glutamate (Log_2_FC: -1.99) and L-arginine (Log_2_FC: 2.12), carbohydrates including glucose (Log_2_FC: -3.56) and D-glucosamine (Log_2_FC: -3.84), and nucleotides such as uracil (Log_2_FC: -3.77) and inosine (Log2FC: 5.17). However, as mentioned above, many of these metabolites are included in commercially formulated culture media, and are therefore unlikely to influence cell viability. Interestingly, whilst lipids are often poorly detected on LC-MS metabolomics platforms using HILIC chromatography where most non-polar species elute with little or no column retention and are thus subject to substantial ion suppression effects in mass spectrometry, many peaks putatively annotated as phosphatidylethanolamines (PEs), and in addition several phosphatidylcholines (PCs) and phosphatidylinositols (PIs), were detected only in goat serum (S1 Table). Furthermore, L-carnitine and O-acetylcarnitine were shown to be significantly lower in FBS relative to goat serum (Table 1). Altogether, many of the putatively identified metabolites significantly altered between GS and FBS appeared to be lipid-like in nature (S1 Table). This indicated that lipids may be a key influencer of cell culture viability, as this metabolite class is only provided to cells in serum supplements.

**Figure 2:**
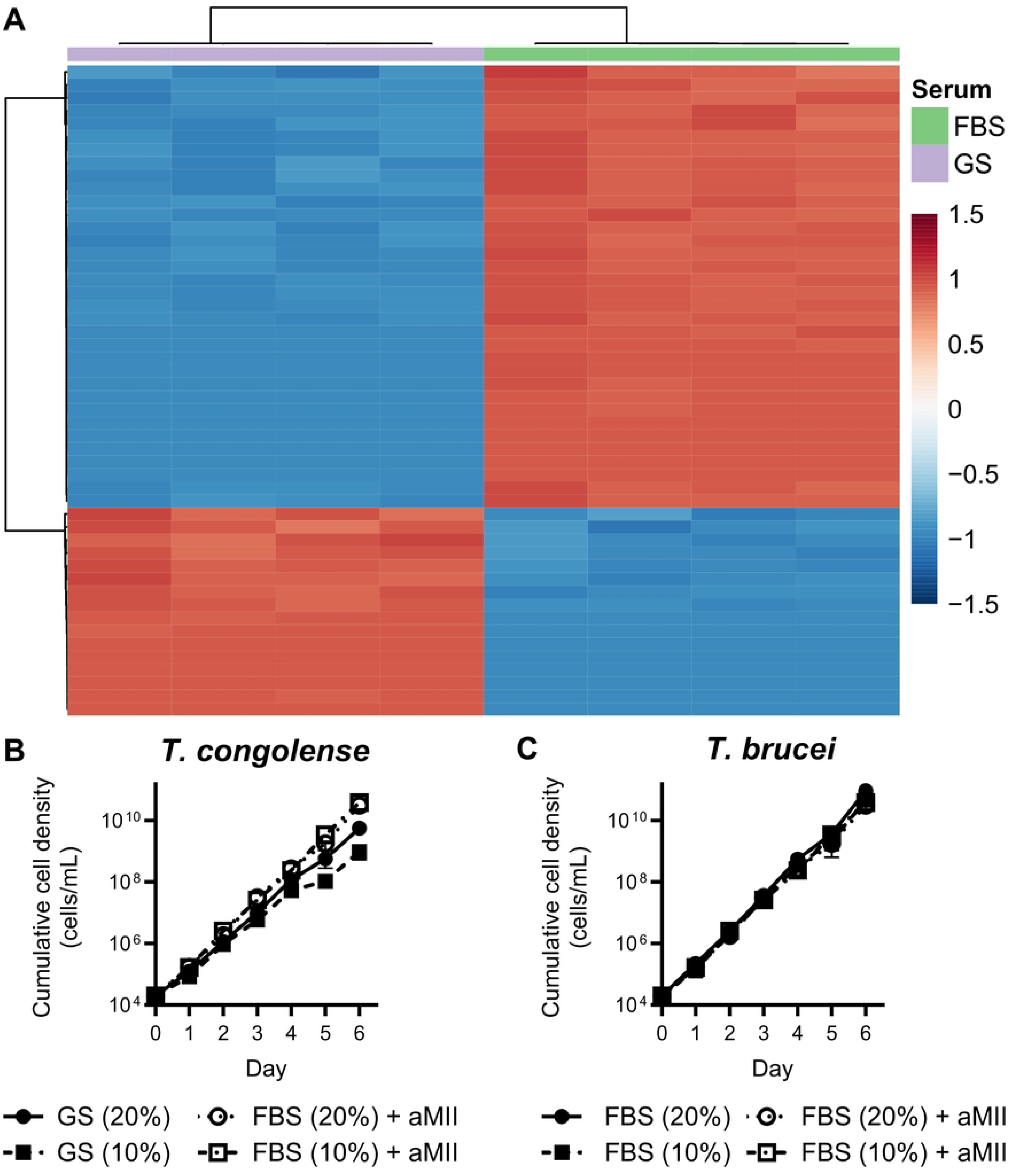
Adaptation of *T. brucei* and *T. congolense* to a universal lipid-loaded medium. A) LC-MS metabolomics analysis of GS (lilac) and FBS (green) was carried out and fold changes as well as t-tests calculated for each detected metabolite. The 50 most significant differences are shown, highlighting widespread differences in polar metabolites between the two sera. B) Cell viability of T. congolense in FBS-supplemented culture with the addition of 2.5% (AMII). Regardless of percentage (v/v) serum, AMII drastically improved cell growth, even compared to the goat serum control. C) Addition of AMII does not impact *T. brucei* growth compared to the FBS controls.

**Table 1:** Biologically relevant metabolites with significant variation in abundance between FBS and GS. Log_2_ fold-change (FC) analysis (GS/FBS) is summarised alongside their associated P-values (*t*-test) and false discovery rate (FDR)-adjusted P-values. Positive values indicate higher abundance in GS, negative values indicate higher abundance in FBS. The full dataset is available in the supplementary results (S1 Table).

| Metabolite | Log<sub>2</sub>FC | P-value | FDR |
| --- | --- | --- | --- |
| Cholate | 19.78 | 5.49E-07 | 3.83E-06 |
| Inosine | 5.17 | 4.77E-06 | 2.24E-05 |
| 3-(4-Hydroxyphenyl)pyruvate | 4.06 | 3.88E-02 | 6.26E-02 |
| L-Carnitine | 2.26 | 8.07E-07 | 5.19E-06 |
| O-Acetylcarnitine | 2.14 | 1.99E-06 | 1.08E-05 |
| L-Arginine | 2.12 | 8.33E-06 | 3.40E-05 |
| Pyruvate | 1.43 | 1.56E-03 | 3.57E-03 |
| Imidazole-4-acetate | 1.42 | 2.42E-05 | 8.53E-05 |
| L-Citrulline | 1.28 | 9.56E-04 | 2.30E-03 |
| L-Asparagine | 1.24 | 2.37E-03 | 5.20E-03 |
| L-Phenylalanine | -1.05 | 7.07E-06 | 2.99E-05 |
| Hypoxanthine | -1.07 | 1.10E-04 | 3.26E-04 |
| Citrate | -1.09 | 5.32E-05 | 1.69E-04 |
| L-Glutamine | -1.09 | 3.21E-05 | 1.10E-04 |
| Adenine | -1.33 | 8.20E-04 | 2.00E-03 |
| Deoxyuridine | -1.69 | 1.48E-06 | 8.60E-06 |
| (R)-Lactate | -1.90 | 1.14E-09 | 7.24E-08 |
| D-Glucose | -3.56 | 1.02E-07 | 1.22E-06 |
| Uracil | -3.77 | 5.17E-10 | 4.99E-08 |
| Succinate | -4.95 | 5.94E-08 | 8.68E-07 |

To probe this further, viability of both species was assessed when grown in FBS with a lipid-loaded BSA supplement (albuMAX II, AMII; 2.5% v/v; Fig 2B & 2C). In the presence of both 20% and 10% FBS supplemented with AMII, *T. congolense* grew at rates comparable to that of the goat serum control (mean doubling time of 10.03 hr, 9.5 hr and 9.1 hr for 20% FBS, 10% FBS and 20% GS, respectively), suggesting the addition of the BSA supplement was key to growth of this species in FBS-supplemented medium *in vitro* (Fig 2B). Cell viability of *T. brucei* was not impacted by lipid supplementation, indicating that a medium formulation consisting of 10% FBS and 2.5% AMII was sufficient for the culture of both trypanosome species (Fig 2C).

### Comparative analysis of the *T. brucei* and *T. congolense* lipidomes

Serum supplements are, typically, the single source of exogenous lipid precursors for *in vitro* culture systems. Therefore, the apparent requirement for exogenous lipids in *T. congolense* may indicate differences in how this species obtains and utilises the host’s lipids it requires for cell survival compared to *T. brucei*. This suggests that *T. congolense* preferentially scavenges lipids from its host, rather than synthesising *de novo* the required lipid complement.

Interrogation of the *T. brucei* and *T. congolense* genomes suggests that *T. congolense* shares the majority of enzymes linked to lipid metabolism, including all four fatty acid elongases (ELO1-4; S2 Table), as previously described [8]. However, no orthologue was found in either of two independent *T. congolense* genome assemblies (shotgun or PacBio, released in 2016 and 2019, respectively) for the two *T. brucei* genes encoding putative fatty acid desaturases, Tb927.10.7100 and Tb11.v5.0580. The protein encoded by Tb11.v5.0580 was recently found to desaturate long chain FAs (20:4 and 22:4) resulting in the generation of docosapentaenoic acid (22:4) and docosahexaenoic acid (22:6), and has been hypothesised to play a role in BSF differentiation to stumpy-form [12]. However, it is difficult to speculate on precise lipid metabolic distinctions using genomic information alone, particularly as many of these predicted enzymes remain to be experimentally validated in either trypanosome species.

In order to compare differences in lipid composition further, untargeted LC-MS lipidomics analysis was performed on both *T. brucei* and *T. congolense* BSFs, with *T. congolense* grown in HMI-93 with goat serum as well as HMI-11 with 10% FBS and 2.5% albuMAX II, in order to directly compare parasite lipidomes when grown under identical conditions (with one caveat that *T. brucei* is grown at 37°C, and *T. congolense* at 34°C). Even under these circumstances, *T. brucei* and *T. congolense* bloodstream forms displayed distinct lipidomic profiles (Figure 3).

**Figure 3:**
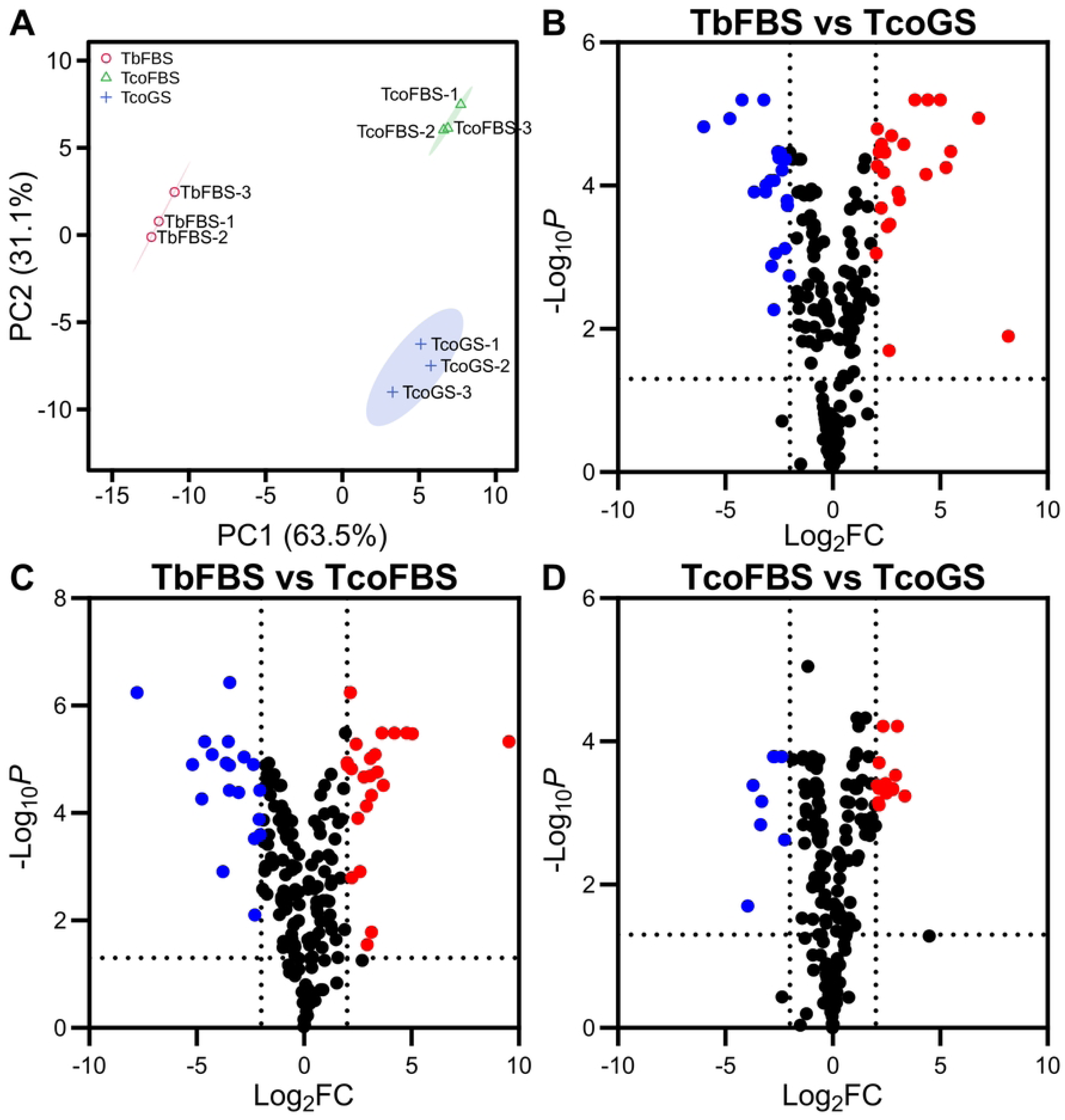
Overview of comparative LC-MS lipidomics analysis of *T. brucei* and *T. congolense*. A) PCA plot of three sample groups: TbFBS, TcoGS and TcoFBS. The first and second principal components are plotted with 95% confidence regions shown, highlighting separation of species (PC1) and culture conditions (PC2). Data were Log_2_ transformed and auto-scaled prior to PCA loading. B) Volcano plot showing significantly altered (*p* < 0.05 & Log_2_FC > 2 [Red], < -2 [Blue]) lipid species when comparing *T. brucei* culture in FBS and *T. congolense* cultured in GS. A total of 46 lipids were significantly altered under these criteria. C) Volcano plot highlighting the 42 lipid species that were significantly altered when comparing *T. brucei* and *T. congolense* grown under the same in vitro conditions (10% FBS, 2.5% AMII). D) Volcano plot showing the 22 lipid species that were significantly altered when comparing *T. congolense* cultured in GS and FBS, 2.5% albuMAX II.

LC-MS lipidomics analysis resulted in the identification of 209 lipids across the three sample groups (Fig 3; S3 Table). PCA analysis showed clear separation of the three sample groups, with PC1 (explaining 63.5% of variation) separating primarily by species, whilst PC2 (explaining 31.1% of variation) separated primarily by serum content (Fig 3A). Statistical analyses (BH-corrected *t*-tests) were applied to three comparisons - TbFBS vs TcoFBS, TbFBS vs TcoGS and TcoFBS vs TcoGS - highlighting a total of 46, 42 and 22 significantly altered (Log_2_ fold change [FC] >2 & *p* < 0.05; Fig 3B-D) lipid species in the respective comparisons. Notably, there were no clear patterns in the 11 lipid species that were significantly altered between the two *T. congolense* sample groups, suggesting that the switch from goat serum to FBS/AMII did not lead to widespread lipidomic changes (Fig 3D).

Given the data was normalised to cell number, within-feature comparisons and patterns across groups were analysed in more detail (rather than absolute quantification), with a particular focus on lipids significantly altered in the aforementioned two-way analyses. The most abundant group of lipids present across the sample groups were glycerophospholipids (phosphatidylcholines, PCs; phosphatidylethanolamines, PEs; *lyso*-phosphatidylcholines, LPCs; phosphatidylinositols, PIs; 61.7% of identified lipids, S3 Table).

PCs, in particular short chain PCs, were consistently more abundant in *T. brucei* (S3 Table), with a comparative reduction in *T. congolense* (for both FBS and GS sample groups; Fig 4A). In contrast, for all significantly altered PCs recovered, none exhibited differences between the two *T. congolense* sample groups. Notably, longer chain PCs with higher levels of unsaturation (e.g. PC 36:6, 40:8 and 40:9) were more abundant in *T. congolense*, with a switch to FBS leading to reduced levels of PC 40:9 in particular (Fig 4A). These data indicate that *T. congolense* exhibits a preference for longer acyl chain PCs.

**Figure 4:**
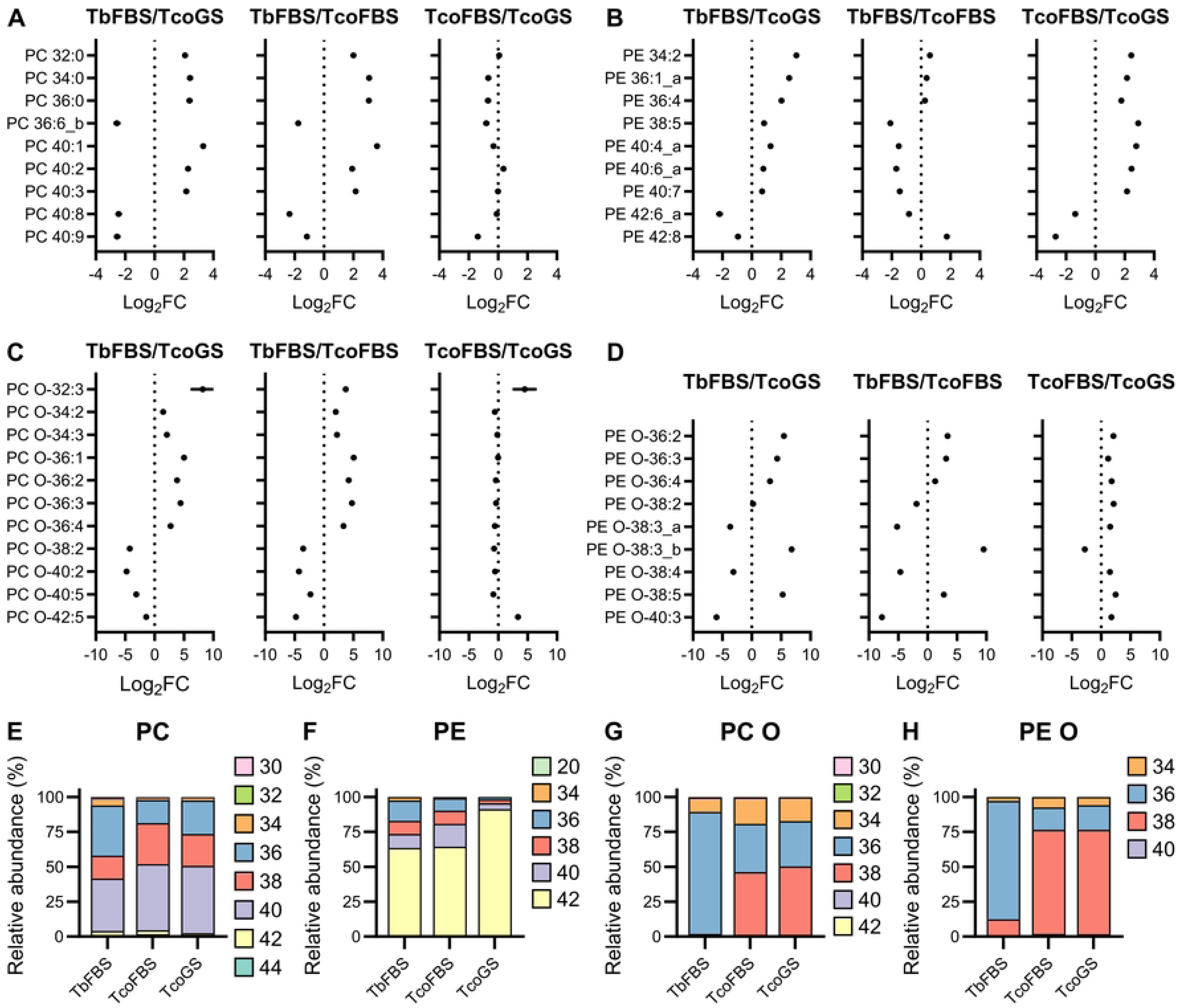
PC and PE profiles in *T. brucei* and *T. congolense*. Forest plots were generated for each lipid classes to show fold changes of lipids that were significantly altered in one of the two-way comparisons. A) Nine PCs were significantly altered, with *T. brucei* showing elevated abundance of 6 PCs compared to *T. congolense* in both conditions. PCs did not vary between the two *T. congolense* sample groups, regardless of serum supplement. B) There were also 9 PEs significantly altered between the three sample groups. In this case, shorter PEs were generally more abundant in cells grown in FBs, regardless of species. C) A total of 11 PC-Os were significantly altered, in each case the significant change in abundance was a species-specific difference. Shorter chain PC-Os were generally more abundant in *T. brucei*, whilst longer chain PC-Os were more abundant in *T. congolense*. D) Nine PE-O lipids were significantly altered in this data. Similar to PC-Os, the PE-Os showed species-specific differences in abundance, with very little variation between the two *T. congolense* sample groups. Where multiple isomers were detected, these are indicated with an underscore and letter. E) Stacked bar plot showing each carbon chain length as a percentage of the total PC pool, highlighting a reduction in 36-carbon chain lipids in *T. congolense*. F) A stacked bar plot showing carbon chain length abundances for PEs, this time highlighting a serum-dependent change, with *T. congolense* cultured in GS exhibiting almost exclusively 42-carbon length lipids. G) A stacked bar plot for PC-O, highlighting a species-specific difference, with *T. congolense* exhibiting a substantial increase in 38-carbon chain length PC-O lipids. H) PE-O lipids exhibited a very similar species-specific different to PC-Os, with T. congolense exhibiting large amounts of 38-carbon lipids in this pool.

In general, abundances of significantly altered PEs were higher in *T. brucei*, regardless of chain length (Fig 4B). Notably, *T. congolense* cultured in GS-supplemented media exhibited lower abundance of almost all PEs identified, compared to both FBS sample groups. This was with the exception of long chain unsaturated PEs, in particular PE 42:6 and 42:8 (Fig 4B), mirroring the PC lipid class that suggested a preference for longer acyl chain lipids for *T. congolense*.

The most prominent species-related difference in these data was found in ether-linked glycerophospholipid abundance (PC-O and PE-O; Fig 4C & 4D, respectively). For both lipid classes, *T. congolense* exhibited significantly reduced abundance compared to *T. brucei* (Fig S2A & S2B), regardless of culturing conditions. A pattern was observed here, whereby *T. brucei* exhibited significantly increased abundance of shorter chain PE-Os and PC-Os (e.g. PE-O 36:x and PC-O 36:x; Fig 4C & 4D, respectively), whilst both *T. congolense* samples showed elevated levels of longer-chain lipids in both classes, in particular PC-O 38:2, 40:x and 42:5 (Fig 4C; shown on a logarithmic scale in Fig S2C) and PE-O 40:3 (Fig 4D). PC-O and PE-O lipid abundances were only significantly altered in inter-species comparisons, with the two *T. congolense* samples exhibiting very similar profiles for ether-linked glycerophospholipids (Fig 4C & 4D). Notably, *T. brucei* exhibited elevated abundance of a PC-O 38:3 putatively annotated as PE(O-20:0/18:3) (38:3_b), whilst an isomer annotated as PE(O-18:0/20:3) (38:3_a) was elevated in *T. congolense* (Fig 4D), indicating differences in lipid remodelling between the two species. Finally, *T. congolense* cultured in GS exhibited a high abundance of several LPCs (Fig S2A), when compared to either species cultured in FBS. In particular, ether-linked LPCs (20:0, 20:1 and 20:2) were significantly elevated in abundance (Fig S2D). Notably, the total LPC content in each sample group remained below 0.4% of the total PC pool (S3 Table), confirming that elevated abundances observed in *T. congolense* GS samples reflects a true biological difference instead of an artifact resulting from lipase activity during lipid extractions.

To further determine whether *T. congolense* does indeed exhibit a preference for longer chain lipids compared to *T. brucei*, abundances of each carbon chain length were summed (regardless of chain saturation) for each lipid class and graphed as a percentage of the total abundance of that lipid class (Fig 4E, 4F, 4G & 4H). Whilst the PC pool in *T. brucei* was >40% 36-chain lengths, this was reduced in *T. congolense* under both GS and FBS-supplemented conditions, with *T. congolense* instead exhibiting increased abundance of 38 and 40-carbon chain lipids (Fig 4E). The PE pools of *T. congolense* and *T. brucei* were strikingly similar under FBS-supplemented conditions, dominated by 42-carbon chain lipids (∼60%). In contrast, when cultured with GS-supplementation, the *T. congolense* PE pool was almost exclusively 42-carbon chain length (Fig 4F). For both PC-O (Fig 4G) and PE-O (Fig 4H), a strong species-specific difference was observed, with *T. congolense* exhibiting a high abundance of longer chain lipids (in particular 38-carbon chains), regardless of culturing conditions.

There were fewer clear species- or condition-specific patterns in other lipid classes, in particular sphingolipids (sphingomyelins, SM) and PIs. For lipids that were significantly altered in two-way comparisons, these were generally higher in abundance in *T. congolense*, especially when parasites were cultured in the same medium (Fig 5A). Similar to above, longer chain SMs (42:x and 44:2) were highly abundant in *T. congolense* (Fig 5A), whereas the *T. brucei* SM pool was almost exclusively made up of 40-carbon chain SMs (Fig 5B).

**Figure 5:**
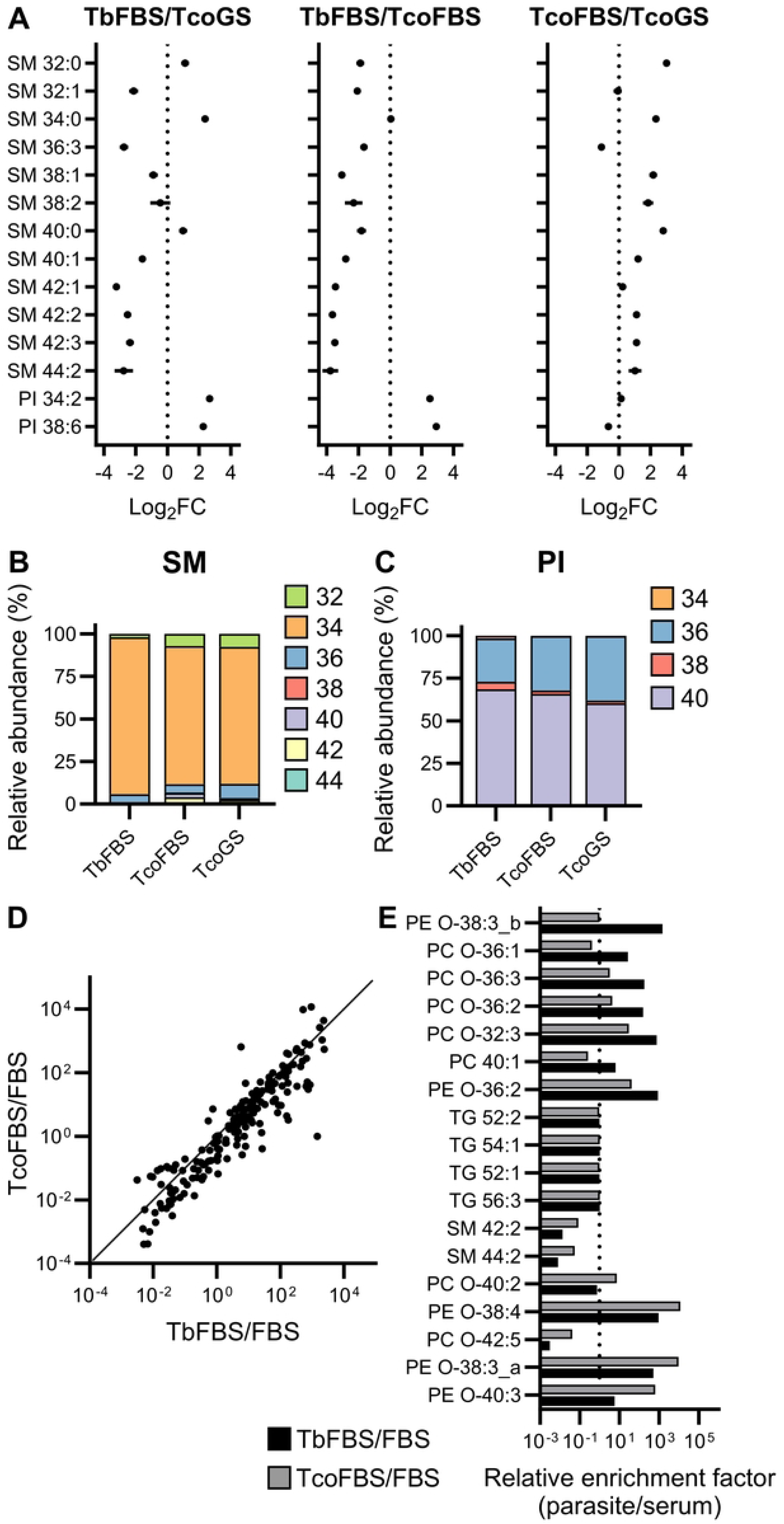
PI and SM profiles, and lipid uptake ratios in *T. brucei* and *T. congolense*. A) Forest plot to show abundance changes in 12 SMs and 2 PIs that were significantly altered in at least one of the two-way comparisons. In general, these data highlighted a reduction of longer-chain SMs in *T. brucei* compared to *T. congolense*. B) Stacked bar plot showing each carbon chain length as a percentage of the total in the SM pool. Unlike glycerophospholipids, differences in the SM pool were not as drastic, although *T. congolense* exhibited a small increase in 32- and 36-carbon chain SM lipids, compared to *T. brucei*. C) Stacked bar plot showing overall chain length composition of the PI pools in *T. brucei* and *T. congolense*. Both species showed highest abundance of 40-carbon chain length lipids in this class, with no significant variation between the three sample groups tested. D) Global XY scatter plot comparing the relative uptake ratios of individual lipid species in *T. brucei* (TbFBS vs FBS) and *T. congolense* (TcoFBS vs FBS). The solid diagonal line represents the line of identity (x = y), indicating equivalent relative lipid accumulation between the two species. While shared lipid pools cluster along this line of identity, divergence is evident in several lipids. E) Targeted relative enrichment factor for 18 representative lipids. A base line of 1.0 (indicated by the vertical dashed line) indicates likely passive equilibration with the host serum, as observed universally for neutral storage triglycerides (TGs). Values deviating from 1.0 highlight species-specific accumulation. For example, *T. brucei* exhibits selective accumulation of PE-O 38:3_b and PC-O 36:x. In contrast, *T. congolense* exhibits selective accumulation of longer chain lipids such as PC-O 40:2 and PE-O 40:3. The full dataset is presented in S4 Table.

The remaining lipid classes identified were neutral lipids, cardiolipins and cholesterol esters (Fig S2B). Two cardiolipins were significantly altered in a species-specific manner, with CL 76:12 being highly abundant in *T. brucei*, and in contrast, CL 76:8 being highly abundant in *T. congolense* under both GS and FBS-supplemented conditions (Fig S2B).

It was recently shown that *T. congolense* is highly resistant to inhibitors of fatty acid synthesis, leading to the hypothesis that this species preferentially scavenges lipids from the exogenous environment. To evaluate this hypothesis, the sera used to culture *T. brucei* and *T. congolense* were also analysed by LC-MS lipidomics (S3 Table). Spearman rank correlations were subsequently performed to compare the parasite lipidomes with their respective sera lipidomes (Fig S3, S4 Table). However, correlations of different lipid classes between the parasites and sera were only moderate to weak (with the exception of storage lipids - MG, DG and TG). This lack of global correlation indicated that neither *T. brucei* nor *T. congolense* scavenge lipids indiscriminately. Therefore, to quantify whether host lipids are selectively scavenged, individual relative uptake ratios (lipid abundance within the parasite relative to the serum) were calculated for each lipid species detected in both the parasites and the matched serum samples, with a focus on FBS (Fig 5D & 5E; S4 Table).

When plotted together, the majority of lipids showed similar relative uptake ratios between *T. brucei* and *T. congolense*, mapping closely to the line of identity (x = y; Fig 5D). This is highlighted by the aforementioned neutral storage lipids, in particular TGs, with uptake ratios for both species of approximately 1.0 (Fig 5E). However, there were several lipids that deviated significantly from the identity line, highlighting species-specific differences in lipid uptake. The majority of lipids exhibiting this divergence were glycerophospholipids (Fig 5E).

To further characterise species-specific lipidomic differences, a previously published comparative transcriptomics dataset [8] was analysed with a focus on lipid enzymes (Fig S4A & S4B). Notably, these data were generated in HMI-11 and HMI-93 (supplemented with 20% goat serum) for *T. brucei* and *T. congolense*, respectively [8]. Firstly, RNAseq data was mapped to glycerophospholipid metabolism (Fig S2C), where a bottleneck was observed at ether-lipid initiation. Transcripts corresponding to dihydroxyacetone phosphate acyltransferase (DAT) were not detected in *T. congolense* samples (S2 Table), and furthermore, transcripts corresponding to alkyl-DHAP synthase (ADP) were reduced in this species, compared to *T. brucei*. This bottleneck could explain the marked reduction in ether-glycerophospholipid abundance in *T. congolense*, although the reasons for lack of activity in the pathway are currently unknown.

Within glycerophospholipid headgroup assembly pathways, there was a reduction in transcript abundance corresponding to PE synthesis in *T. congolense* (ECT & EPT; Fig S2C). In contrast, transcripts corresponding to PC synthesis (CCT & CPT; Fig S2C) exhibited upregulation in this species, compared to *T. brucei*, supporting the lipidomic differences observed, where PE species were generally lower in abundance in both *T. congolense* sample groups (Fig 4). Of further note, transcripts associated with PLA1 were higher in *T. brucei* (Fig S2C), supporting higher LPC production in this species. Increased levels of LPC 22:5 and 22:6 could also be explained by exogenous lipid bioavailability [13,14].

Sphingolipid and long-chain FA metabolism were subsequently analysed (Fig S2D). In general, transcripts corresponding to these pathways were most abundant in *T. brucei*, with key exceptions being ELO4 and ACS5, suggesting species-specific channelling of acyl-CoA into sphingolipid and glycerophospholipid synthesis. These transcriptomic differences could contribute to the distinct chain-length distributions seen in the corresponding lipidomes. Taken together, species-specific differences in previously published transcriptomics data support the species-specific differences observed in lipidomics analyses carried out in this study.

### Comparative analysis of *T. brucei* and *T. congolense* GC-MS fatty acid profiles

Overall, the above results point towards differences in the fatty acyl composition of *T. congolense* lipids relative to *T. brucei*. To examine this further, targeted GC-MS fatty acid analysis was conducted for the three sample groups explored above: *T. brucei* in FBS/AMII, *T. congolense* in GS and *T. congolense* in FBS/AMII (Table 2; raw data in S5 Table). It is known that *T. brucei* BSFs contain high levels of C18:0, C18:1 and C18:2 fatty acids, which is also evident from the current analysis. Additionally, *T. brucei* also display substantial enrichment of C22:4, C22:5 and C22:6 PUFAs relative to their hosts (or cell culture serum) [13], indicating the parasites either have a mechanism for selectively acquiring these fatty acids from their environment or input significant resources to *de novo* biosynthesise these fatty acids. Although BSF *T. congolense* are shown to contain similar fatty acid species to *T. brucei*, their relative abundance differs – the ratio of fatty acids C22:4 and C22:6 increases relative to levels of C18:2, the most abundant fatty acid in both trypanosome species (Table 2). There is a drop in relative cellular levels of C22:6 when *T. congolense* parasites are moved from goat serum to FBS/AMII-based culture, suggesting supplies of this fatty acid are supported by exogenous sources, and that abundance of this PUFA is reduced in the latter supplement. Finally, *T. congolense* exhibited increased abundance of C16:0 compared to T. brucei, irrespective of culturing conditions.

**Table 2:** Comparative GC-MS fatty acid analysis of *T. brucei* and *T. congolense*. Abundance of each FA is shown as a percentage of total detected FA.

| Fatty acid | *T. brucei* (FBS) | *T. congolense* (FBS) | *T. congolense* (GS) |
| --- | --- | --- | --- |
| (C15:0) | 0.00 | 0.10 | 0.25 |
| (C16:0) | 1.37 | 14.44 | 12.86 |
| (C17:0) | 0.39 | 0.65 | 0.72 |
| (C18:0) | 20.78 | 14.52 | 14.62 |
| (C18:1) | 8.55 | 8.50 | 5.58 |
| (C18:2) | 27.43 | 26.06 | 25.94 |
| (C18:3) | 2.78 | 5.39 | 4.88 |
| (C20:0) | 0.00 | 0.34 | 0.03 |
| (C20:2) | 0.00 | 0.10 | 0.28 |
| (C20:3) | 2.24 | 0.96 | 0.00 |
| (C20:4) | 0.62 | 0.85 | 1.04 |
| (C22:4) | 5.88 | 6.13 | 4.30 |
| (C22:5) | 19.82 | 15.33 | 16.28 |
| (C22:6) | 10.13 | 6.63 | 13.22 |

## Discussion

*T. congolense* and *T. brucei* both cause AAT. Whilst closely related and historically assumed to be biologically similar, emerging data suggests that these species differ significantly at the genome [15], transcriptome [16], proteome [17] and metabolome [8] level. Furthermore, this divergence likely underlies important phenotypes relevant to infection biology, disease, and disease control. Whilst *in vitro* culture and genetic manipulation of *T. congolense* has become routine over the past decade, there remain notable and peculiar differences compared to *T. brucei* and, indeed, many other axenic eukaryotic *in vitro* systems. For example, *T. congolense* is only viable with a GS supplement instead of FBS [7]. Secondly, unlike most eukaryotes, *T. congolense* BSF are only viable at 34°C, rather than 37°C [7]. The reasons underlying these peculiarities are currently unknown, but they place limitations on carrying out comparative inter-species analyses under *in vitro* conditions.

In this study, *in vitro* culture of *T. congolense* was optimised by testing various mammalian-derived sera. *T. brucei*, was not viable in CS or ABS, whilst *T. congolense* growth was not viable in FBS, as previously reported. Given that serum is the primary source of lipids, growth in HMI-11 was subsequently tested with the addition of an alternative lipid source - AMII. Interestingly, the addition of AMII to HMI-11 supported *T. congolense* growth to a greater degree than HMI-93 (containing 20% goat serum), indicating that exogenous presence of lipids is important for *T. congolense* viability, although further work is required to understand how this translates to the situation *in vivo*. It is currently unclear whether the apparent lipid requirement of *T. congolense* influences host-adaptation, or restricts host specificity (e.g. *T. congolense* is known to infect many non-ruminants). Importantly, the HMI-11/AMII formulation enables the *in vitro* culture of both *T. brucei* and *T. congolense* in the same medium (this was not previously possible), allowing for their direct comparison.

Given the growth phenotypes observed in this study, a comparative lipidomic characterisation was carried out between the two trypanosome species. Even when grown under the same media conditions these two parasite species display distinct lipidomic profiles. Most lipid metabolic genes that have been identified and functionally characterised in *T. brucei* to date have orthologues in the *T. congolense* genome. Therefore, despite this apparent genomic similarity, functional divergence has occurred between the two species in terms of lipid metabolism. Indeed, previous analyses showed that *T. congolense* is less sensitive to specified inhibitors (e.g. Orlistat & acetyl-CoA synthetase inhibitors) of fatty acid synthesis than *T. brucei*, indicating that *T. congolense* preferentially scavenges lipids from its host environment, *in lieu* of biosynthesis [8].

This study reveals two important aspects of lipid metabolism in trypanosomatids. Firstly, *T. congolense* appears to have a preference for longer chain polyunsaturated lipids, compared to *T. brucei*, with notable enrichment observed among glycerophospholipids (for example, PC 40:8, PC 40:9, PE 42:8, PI 34:2 & PI 38:6). Whilst it could be hypothesised that this is a result of increased scavenging, correlation between the *T. congolense* lipidome and host serum lipids is low, similar to *T. brucei*. Instead, it is likely that *T. congolense* elongates and remodels lipids to a greater extent, compared to *T. brucei*, a mechanism supported by increased transcript abundance of ELO4 and ACS5. Glycerophospholipids, in particular PC and PE, are key components of cell membranes [18], and increased abundance of these long chain PUFAs could result in increased fluidity, flexibility and elasticity of the plasma membrane [19,20]. Further studies are required to determine the biophysical differences between the *T. brucei* and *T. congolense* plasma membranes.

Secondly, *T. congolense* exhibits very low abundance of ether-linked glycerophospholipids, for reasons currently unknown. Finally, even in shared medium, intrinsic species-specific lipidomic differences exist. *T. brucei* retains much larger ether-linked glycerophospholipid pools and the greatest total SM abundance, whereas *T. congolense* produces comparatively fewer ether lipids.

The biosynthesis of ether glycerophospholipids is known to be initiated at the glycosomes of *T. brucei*, where the precursor dihydroxyacetone phosphate (DHAP) is converted to cytoplasmic alkyl-glycerophosphate through a number of enzyme catalysed reactions. This alkyl-glycerophosphate can then pass to the ER for use in glycerophospholipid biosynthesis [21]. Currently, the wider functional significance of the high levels of ether lipids typically observed in *T. brucei* is unknown. The enzyme dihydroxyacetonephosphate acyltransferase (TbDAT) that underpins ether lipid biosynthesis was shown to be dispensable in procyclic forms [22], which would suggest that ether lipids themselves are non-essential, at least in this life cycle stage. RNAi of ethanolamine phosphotransferase (EPT; the enzyme that catalyses ether-type PE production) in procyclic *T. brucei* led to diacyl-type PE becoming dominant, which points to ether lipids being largely derived through *de novo* biosynthesis rather than host lipid scavenging in this species. However, the picture is less clear in bloodstream form trypanosomes. The relative absence of ether lipids in *T. congolense* could indicate an overall decrease in *de novo* lipid biosynthetic activity relative to *T. brucei*. Indeed, transcriptomics data indicating a reduction in both DAT and the subsequent step in ether GPL biosynthesis, alkyl-dihydroxyacetonephosphate synthase, support this hypothesis.

Sphingomyelin is a major sphingolipid in African trypanosomes, functioning in plasma membranes, as well as being involved in the active trafficking of GPI-anchored variant surface glycoproteins [23]. Both the *T. brucei* and *T. congolense* SM pools are dominated by 34:1, albeit with subtle differences in other detected SMs. For example, *T. congolense* exhibits increased SM 32:1, 36:1 and 36:2 abundance, indicating to some extent, a propensity for longer-chain lipids, similar to that observed for PCs. This is supported by transcriptomics data that highlight increased ACS5 and ELO4 transcript abundance in this species, with both enzymes acting on long acyl chain [14,24].

Functionally, these data highlight potential differing membrane strategies. With increasing sphingolipid carbon chain length, it has been shown that membrane thickness increases, and that degrees of their unsaturation lead to increased disorder [25]. This may account for observed decreases in lateral diffusion rates within membranes holding long chain sphingolipids. Interdigitation of sphingolipids between the two membrane leaflets also increases with increased carbon length and unsaturation. Thus, the differing sphingolipid profile presented by *T. congolense* (presence of longer-chain SMs with increased desaturation) may indicate that membrane microdomain structures, such as lipid rafts, differ from those occurring in *T. brucei*. Indeed, although it was beyond the scope of this current work to perform targeted analysis of the *T. congolense* sterol profile, sterol species present in these parasites may also differ, given the well-established co-regulation of sterols and sphingolipids in eukaryotes [26].

Several limitations temper these conclusions. Firstly, *T. congolense* is cultured at 34°C, compared to 37°C for *T. brucei*. Whilst a 3°C reduction in temperature is likely to impact the lipidome - eukaryotic cells are sensitive to minor temperature fluctuations [27], it more likely reflects the tissue tropism of *T. congolense*, which binds to vascular endothelial cells that leads to peripheral sequestration, likely within the dermal and subcutaneous microvasculature where temperatures are significantly lower than the core body temperature [28]. Furthermore, a universal problem with commercial serum supplements is batch-to-batch variation. In this case, differences attributed to goat serum versus FBS may partially reflect batch variation rather than true species-specific serum lipidome profiles. Comprehensive multi-omics characterisation of standard commercial animal sera such as those carried out for human serum [29] would therefore be of great benefit to standardising in vitro lipidomics and metabolomics.

In summary, this study further underscores fundamental biological differences between *T. brucei* and *T. congolense*, specifically regarding lipid metabolism. It confirms that significant divergence exists in the way these species acquire, synthesise and remodel their lipidomes, with *T. congolense* exhibiting a distinct preference for very long-chain PUFAs. While the functional implications of these findings remain to be fully elucidated, they likely provide a foundation for key aspects of *T. congolense* biology, such as intravascular cytoadhesion. Ultimately, these species-specific lipid profiles likely underlie phenotypic divergence in the *in vivo* biology and in host-pathogen interactions of these two parasite species.

## Materials and Methods

### Cell culture

Bloodstream form (BSF) *T. brucei* strain Lister 427 were routinely maintained in HMI-11 [9] supplemented with 10% FBS, and incubated at 37°C, 5% CO_2_. BSF *T. congolense* strain IL3000 were routinely maintained in HMI-93 [7] supplemented with 20% goat serum, and incubated at 34°C, 5% CO_2_. For comparative lipidomic analysis, both species were maintained in HMI-11 supplemented with 10% FBS and 2.5% w/v AlbuMAX II (Thermo Scientific) at their respective optimal incubation conditions for 1 week (2-3 passages) prior to lipid extraction.

For growth curves, cells were seeded at a seeding density of 2 × 10^4^ cells/mL or 5 x 10^4^ cells/mL for *T. brucei* or *T. congolense*, respectively, in 2 mL cultures in a 24-well plate. Cells were counted daily using a haemocytometer and passaged every two days, or when cell density exceeded 1 × 10^6^ cells/mL.

Various serum supplements were used in this study: Goat serum (GS; Gibco, Cat #: 16210072, Lot #: 2426173), Foetal Bovine Serum (FBS; Gibco, Cat #: 10270106, Lot #: B2915971RP), Adult Bovine Serum (ABS, Capricorn, Cat #: ABS-1B, Lot #: CP20-3340), Horse Serum (HS; Capricorn, Cat #: HOS-1A, Lot #: CP22-5442) and Calf Serum (CS; Capricorn, Cat #: CS-1A, Lot #: 2534176).

### LC-MS analysis of serum

LC-MS analysis was carried out as previously described [8]. Hydrophilic interaction liquid chromatography (HILIC) was carried out by Glasgow Polyomics (Glasgow, UK), using a Dionex UltiMate 3000 RSLC system (Thermo Fischer Scientific) coupled to a ZIC-pHILIC column (150 mm × 4.6 mm, 5 µm column, Merch Sequant). The column was maintained at 30°C and samples were eluted with a linear gradient (20 mM ammonium carbonate in water and acetonitrile) over 26 minutes with a flow rate of 0.3 mL/minute.

Sample injection volume was 10 µL and samples were maintained at 4°C before injection. A Thermo Orbitrap Exactive (Thermo Fischer Scientific) was used to generate mass spectra, and was operated in polarity switching mode with the following settings: Resolution: 50,000; AGC: 106; m/z range: 70–1,400; sheath gas: 40; auxiliary gas: 5; sweep gas: 1; probe temperature: 150°C; capillary temperature: 275°C. Samples were run in both positive and negative polarity with the following ionisation: source voltage +4.5 kV, capillary voltage +50 V, tube voltage +70 kV and skimmer voltage +20 V for positive mode; source voltage -3.5 kV, capillary voltage -50 V, tube voltage -70 V and skimmer voltage -20 V for negative mode. Mass calibration was performed for each polarity immediately prior to each analysis batch. The calibration mass range was extended to cover small metabolites by inclusion of low-mass contaminants with the standard Thermo calmix masses (below *m/z* 1400), C_2_H_6_NO_2_ for positive ion electrospray ionisation (PIESI) mode (*m/z* 76.0393) and C_3_H_5_O_3_ for negative ion electrospray ionisation (NIESI) mode (*m/z* 89.0244). To enhance calibration stability, lock-mass correction was also applied to each analytical run using these ubiquitous low-mass contaminants. A set of authentic standards was run prior to the sample set.

RAW spectra were converted to mzXML files (mzML files for fragmentation data) using XCMS for untargeted peak detection [30]. The resultant files were further processed using mzMatch [31] for peak matching and annotation, resulting in a tabular output that was analysed using IDEOM with default settings [32]. Statistical analyses were carried out using MetaboAnalyst (v6.0) [33].

### Lipid extraction

Lipid extracts were prepared using the Folch extraction method [34]. 1 x 10^8^ cells were rapidly quenched using a dry ice/EtOH bath, then centrifuged at 1300 x g, 10 min, 4 °C. Cell pellets were resuspended in a minimal volume of media and transferred to 1.5 ml eppendorf tubes, and were centrifuged at 2500 x g, 5 min, 4 °C. The pellets were then washed 1 x 1 ml PBS (centrifugation 2500 x g, 5 min, 4 °C). Pellets were resuspended in 50 ul PBS, and transferred to a glass vial containing 800 ul 2:1 CHCl_3_:MeOH. The vial was centrifuged, then 150 ul H_2_O was added to induce biphasic partition. Vials were then vortex mixed for 1 hr at 4 °C. Samples were centrifuged at 1000 x g, 10 min and the organic (lipid-enriched phase) was transferred to a fresh glass vials. 10 ul of each extract was contributed to a pooled QC vial, for use in downstream mass spectrometry (MS) analysis. Samples were dried under nitrogen and stored at -80 °C prior to analysis. For cell culture serum samples, extraction procedure was as described above except 100 ul of serum was mixed directly with 800 ul 2:1 CHCl_3_:MeOH and 100 ul H_2_O.

### LC-MS lipidomic analysis

Immediately prior to analysis, dried samples were resuspended in 500 ul MeOH. 100 ul of each sample was then mixed with 100 ul MeOH 5 mM ammonium formate, and samples were centrifuged at 11,000 x g, 5 min. The supernatant (∼ 200 ul) was then transferred to an instrument sample vial.

Untargeted lipidomics analysis was performed using a binary HPLC (Accela, Thermo Fisher Scientific) coupled to an electron spray ionization (ESI) and orbitrap mass analyser (Exactive, Thermo Fisher Scientific). Separation was conducted using a Hypersyl GOLD C18 column, 100×2.1 mm 1.9nm particle size (Thermo Fisher Scientific) maintained at 50 °C. Lipids were eluted over 25 minutes at constant flow rate 400 μL min^-1^ using the solvent system A – H_2_O 10 mM ammonium formate, 20 mM formic acid; B – 9:1 isopropanol:acetonitrile 10 mM ammonium formate 20 mM formic acid, with gradient: 65% A, 35% B (0.5 min); 35% A, 65% B (4 min); 0% A, 100% B (19 min); 65% A, 35% B (21.1 min). Water and acetonitrile were HPLC grade and obtained from Thermo Fisher Scientific, IPA was LC-MS grade (Hypergrade LiChrosolv, Merck). Ammonium formate and formic acid were both LC-MS grade and obtained from Sigma Aldrich. Mass spectra were acquired in both positive (POS) and negative (NEG)

Raw data was imported into Progenesis QI v2.4 (Non-Linear Dynamics), where automatic alignment and peak picking was performed. Relative fold quantification was performed by the software using all ion normalization, followed by data filtering based on ion intensity (>10 000), ANOVA score (< 0.05), fold change (> 2) and a coefficient of variance threshold of 30%. Lipidmaps and HMDB database searches were performed to assign lipid annotations. Lipid annotations were assigned based on expected retention time characteristics, mass error thresholds (< 5 ppm), isotope similarity (>80) and expected adduct formation, with reference to retention times and adduct formations observed for a lipid standard mix used to monitor instrument mass calibration. This was performed for both positive and negative ionisation modes.

Tab-delimited output from Progenesis was further filtered manually, and then analysed using MetaboAnalyst (v6.0) [33]. Briefly, missing values were estimated using left-censored data estimated, a low variance filter (interquartile range) was then applied, and data were log_2_ transformed and auto-scaled prior to statistical analyses.

Where two peaks were detected and annotation as the same lipid species, pearson correlation was applied. When r > 0.9, peaks were summed and reported as one lipid; when r < 0.9 lipids were individually reported as “a” and “b”.

### Fatty acid methyl ester (FAME) preparation

Dried total lipid extract, prepared using the Folch method from 1 x 10^8^ cells, was resuspended in 850 ul methanol and 150 ul MeOH:H_2_O (85:15) 8% (v/v) HCl. Samples were then heated at 65 °C overnight. Dried samples were then resuspended in 1 ml 1:1 H_2_O:Hexane and the organic layer containing FAMEs was transferred to a new glass vial. Samples were dried under nitrogen and stored at -80 °C prior to analysis.

### GC-MS fatty acid analysis

FAMEs samples were analysed via gas chromatography mass spectrometry (GC-MS). Gas chromatography was carried out on a Rtx-2330 60 m length × 0.25 mm inner diameter × 0.20 µm film thickness column (Restek, P/N: 10726) installed in a Trace Ultra gas chromatograph (Thermo Fisher Scientific). Helium carrier gas at a flow rate of 1.4 mL/min was used. 1µL of derivatized sample was injected into a Programmable Temperature Vaporiser (PTV) injector using a 1 in 10 split injection. Injection temperature was 80 °C before increasing to a transfer temperature of 220 °C at a ramp rate of 14.5 °C/min. Oven temperature programme was as follows: 80 °C (3.16 min hold); ramp rate 30 °C min^-1^ to 175 °C; ramp rate 3 °C min^-1^ to 210°C; 30 °C min^-1^ to 265 °C (10 min hold). Eluting peaks were transferred through an auxiliary transfer temperature of 200 °C into a ITQ900-GC mass spectrometer (Thermo Scientific) with scanning mass range set to 60-400 m/z. Targeted analysis with comparison to compound FAMEs standards peak areas were extracted using TraceFinder v3.3. Standards and unknowns were determined by using custom R scripts using XCMS 3.8.2, for peak picking, CAMERA peak grouping package, version 1.42.0 and metaMS 1.22.0.

## Acknowledgements

E.D. was supported by a Scottish Metabolomics Network Training grant, a Wellcome ISSF feasibility grant, and funding from the Scottish Universities Life Sciences Alliance. P.S. is supported by a BBSRC Discovery Fellowship (BB/X009807/7). This study was partially supported by a BBSRC grant awarded to L.M. and M.B. (BB/S00243X/1, BB/S001034/1). The Roslin Institute is supported through core funding from the BBSRC (BS/E/D/20002173; BBS/E/RL/230002C). The authors thank the Mass Spectrometry Facility within the MVLS Shared Research Facilities (University of Glasgow) for provision of metabolomics and lipidomics data, and support & assistance.

---

**Figure S1:**
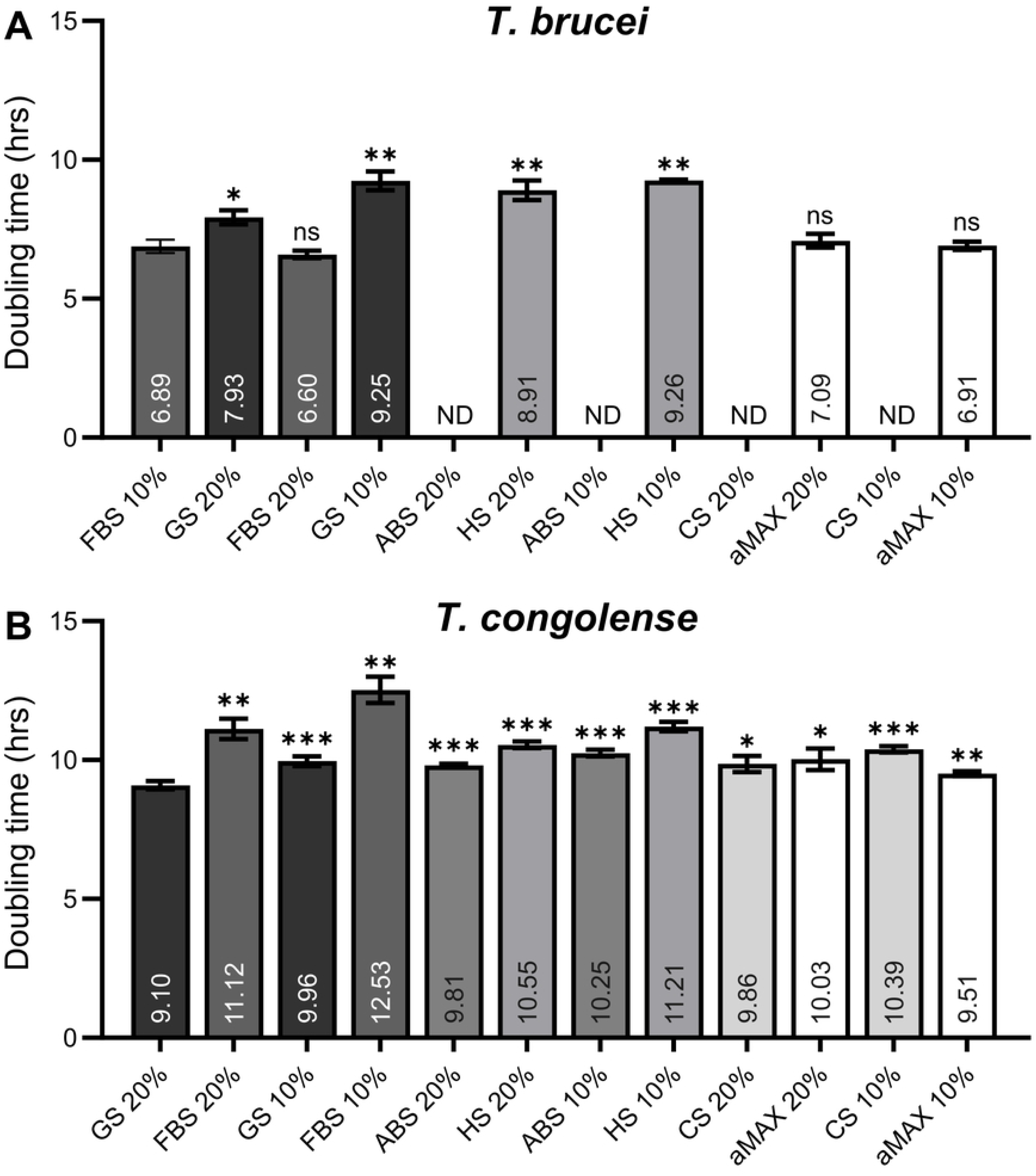
Trypanosome doubling times when cultured in different sera. A) Doubling times for *T. brucei*. B) Doubling times for *T. congolense*. In both panels, mean doubling time is shown (in hrs) inside the bars. Statistical difference was calculated for each condition compared to the control (10% FBS and 20% GS for *T. brucei* and *T. congolense*, respectively) via a BH-corrected *t*-test. Statistics: ns = not significant; * = < 0.05; ** = < 0.01; *** = < 0.001. Abbreviations: ND, not determined; FBS, foetal bovine serum; GS, goat serum; ABS, adult bovine serum; HS, horse serum; CS, calf serum, aMAX, albuMAX II lipid-loaded BSA.

**Figure S2:**
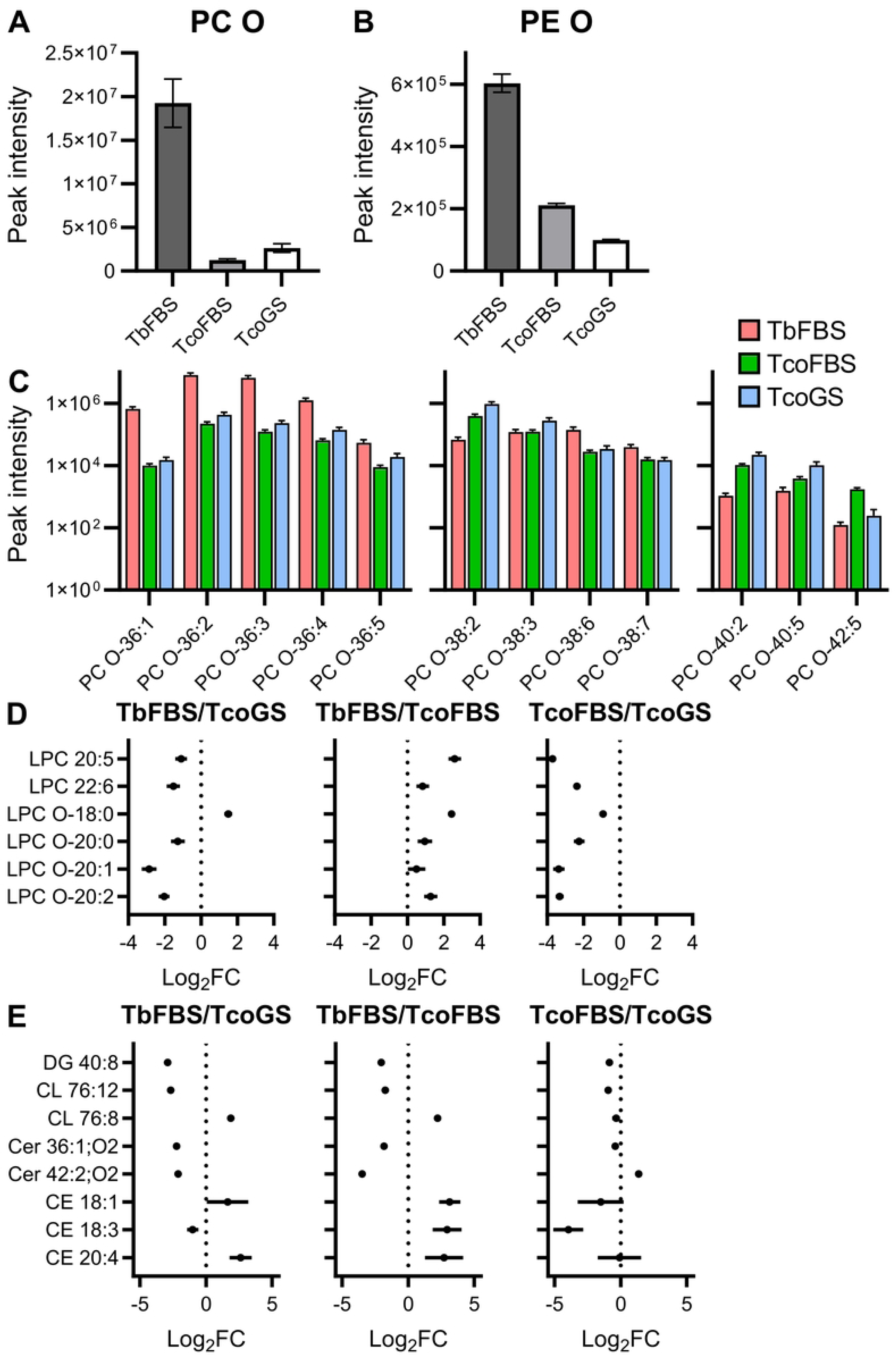
Analysis of lipidomics data. A) Overall abundance of ether-linked PCs (PC-Os) in *T. brucei* (FBS), *T. congolense* (FBS) and *T. congolense* (GS). Overal abundance was calculated by summing the peak intensities of all detected PC-Os, highlighting a significant reduction in both *T. congolense* sample groups. B) Similarly, *T. congolense* exhibits a significant reduction in ether-linked PEs (PE-Os) compared to *T. brucei*. C) *T. brucei* exhibits higher abundance of PC-O 36 lipids, compared to *T. congolense*, regardless of culturing conditions of the latter (left panel). Notably, the degree of increased abundance reduces with increasing desaturation of the 36-carbon chain. Contrastingly, *T. congolense* possesses elevated levels of PC-O 38 lipid species compared to *T. brucei*, although *T. brucei* abundance is higher when the 38-carbon chain is desaturated. The third panel shows 40+ carbon chain length PC-Os, with *T. congolense* exhibiting higher abundance of lipids in this category. D) Forest plots highlighting 6 lysophospholipids that were significantly altered in at least one of the two-way comparisons between the three sample groups. E) Forest plots showing the remaining 8 lipids from various classes that were significantly altered in at least one of the two-way comparisons, including a diglyceride, two cardiolipins, two ceramides and three cholesterol esters.

**Figure S3:**
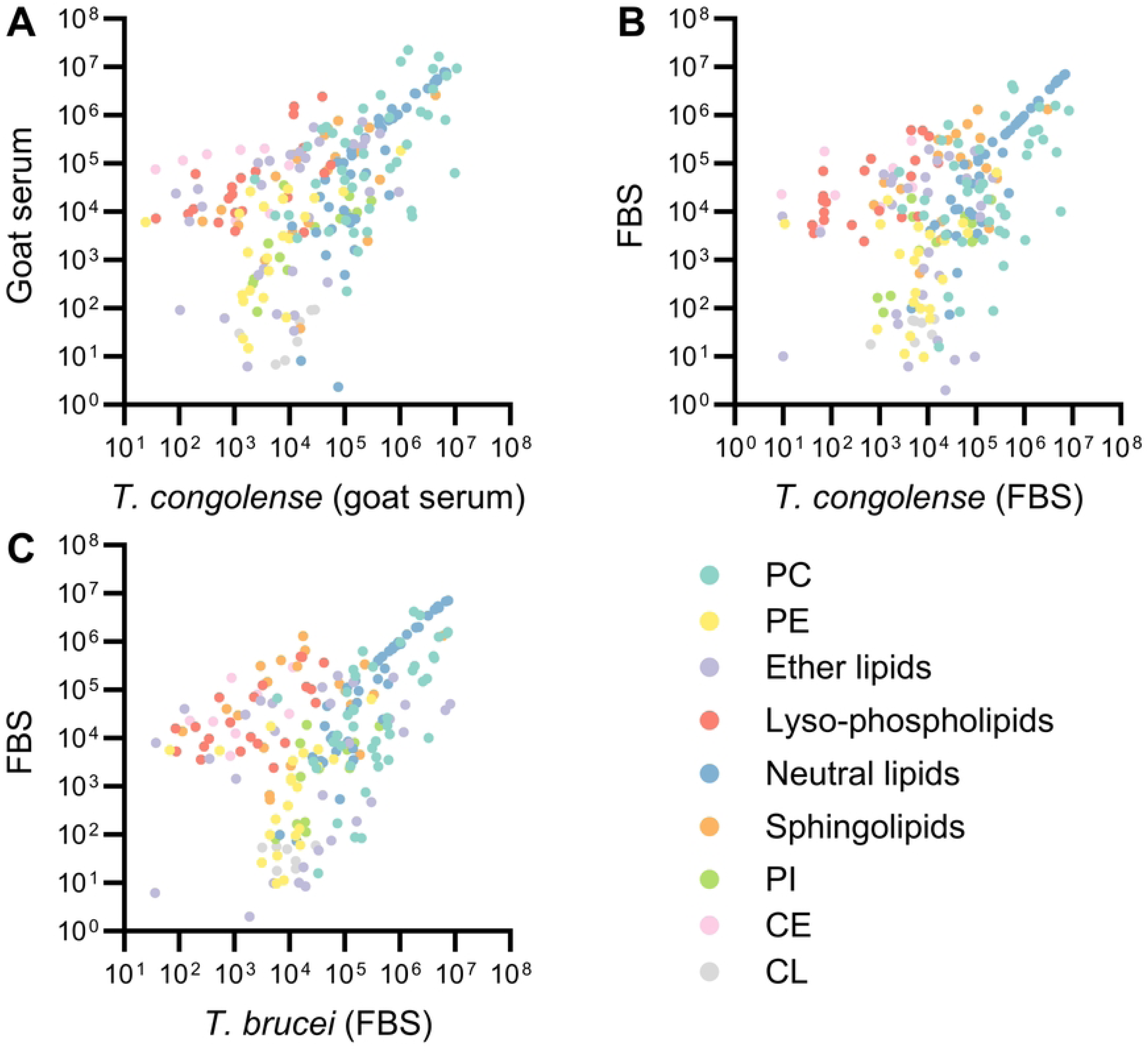
Global correlation analysis of parasite and serum lipidomes. XY scatter plots were generated to display comparisons for (A) TcoGS vs GS, (B) TcoFBS vs FBS, and (C) TbFBS vs FBS. Individual lipid species are colour-coded into 9 lipid classes (indicated in the legend). Only moderate-to-weak correlations were detected (Spearman; S4 Table), with the exception of neutral lipids (>0.9).

**Figure S4:**
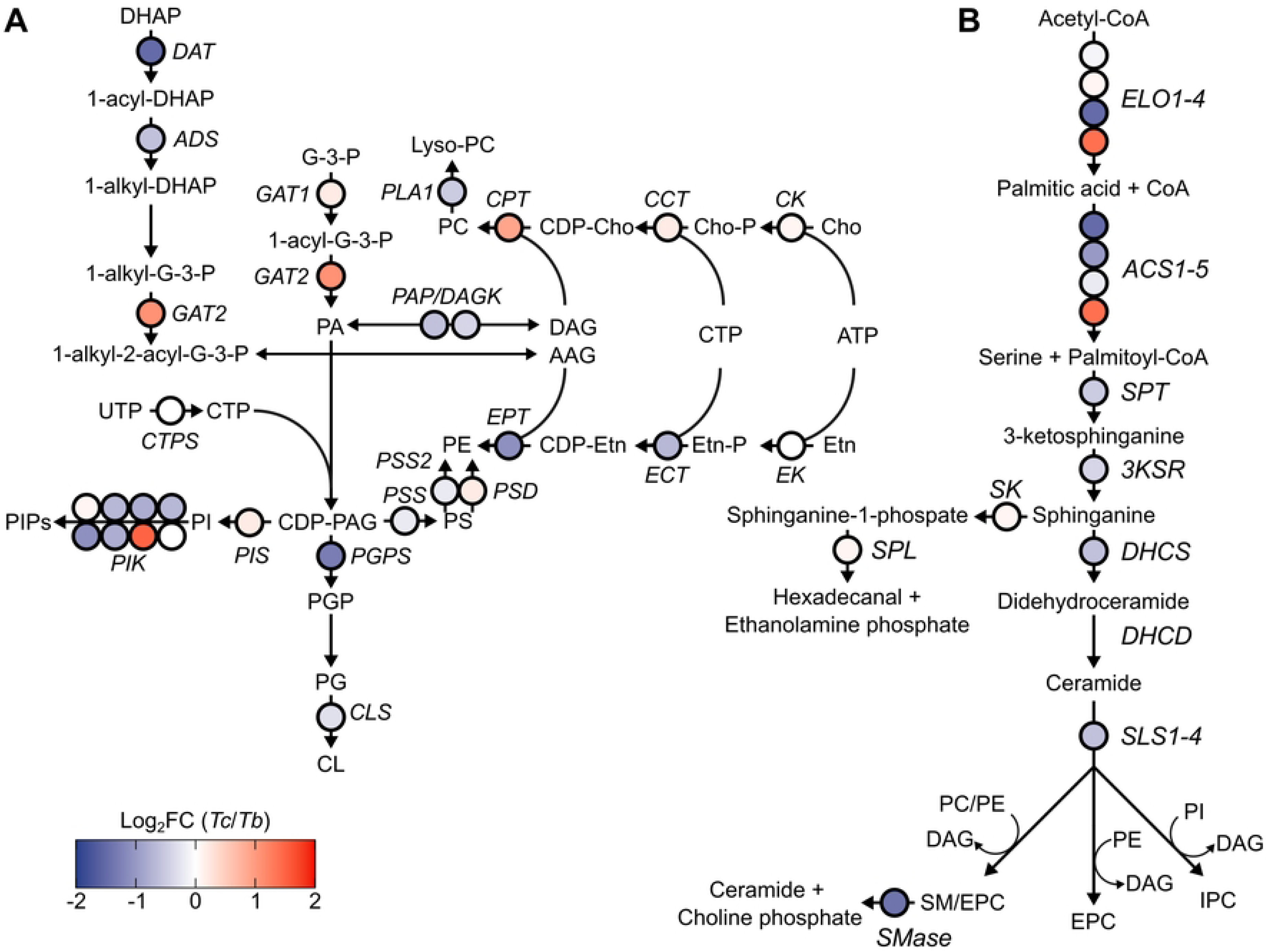
Comparative analysis of lipid metabolism in T. brucei and T. congolense at the transcriptome level. C) RNAseq pathway analysis of glycerophospholipid metabolism. Data were derived from a previous comparative study of *T. brucei* and *T. congolense* [8]. Enzyme names are shown in italics, and dots representing each enzyme are coloured in blue (higher mRNA abundance in *T. brucei*), white (equal abundance) or red (higher mRNA abundance in *T. congolense*). Full enzyme names are listed in S2 table. D) RNAseq pathway analysis of sphingolipid metabolism. Data were derived as mentioned above. Full enzyme names are listed in S2 table.

**S1 Table: Metabolomics analysis of FBS and goat serum.**

**S2 Table: Lipid metabolic enzyme comparison between *T. brucei* and *T. congolense*.** Genes encoding lipid enzymes were previously identified in the TREU 927 reference genome [14]. These genes were used to find orthologues in two *T. congolense* genomes: the shotgun assembly (2016) and the more recent PacBio assembly (2019). These data were combined with transcriptomics data generated previously [8]. This analysis also included an Orthofinder [35] analysis of the *T. brucei* and *T. congolense* proteomes. Where multiple gene IDs are listed in one row, Orthofinder was unable to determine definitively which gene is the true syntenic orthologue, and therefore, the orthologues are grouped together.

**S3 Table: Lipidomics results and analysis.** Microsoft Excel spreadsheet containing all detected and annotated lipids as well as the final dataset used to generate all figures. Separate worksheets are filtered to show only the significant hits obtained via three two-way comparisons: TbFBS vs TcoGS, TbFBS vs TcoFBS and TcoGS vs TcoFBS.

**S4 Table: Parasite and serum lipidome correlation and uptake ratio analysis.** Microsoft Excel spreadsheet containing the Spearman Rank values for 9 lipid classes (PC, PE, ether lipids, lysophospholipids, PI, neutral lipids, sphingolipids, CE & CL) to test for correlations between parasite lipidomes and that of their respective sera. A second worksheet contains uptake ratios for each lipid (parasite abundance vs serum abundance) for TbFBS and TcoFBS compared to FBS.

**S5 Table: GC-MS analysis of *T. brucei* and *T. congolense*.** Raw GC-MS data including retention times and detected masses used to generate Table 2.

